# Genetic depletion of de novo coenzyme A biosynthesis exacerbates puromycin toxicity

**DOI:** 10.1101/2022.09.06.506844

**Authors:** Sunada Khadka, Adam Chatoff, Nathaniel W. Snyder, Ronald DePinho, Florian Muller

## Abstract

Puromycin is an amino nucleoside that inhibits protein synthesis by interrupting elongation of nascent peptide chains. It is a commonly used selection antibiotic in molecular biology research via engineered expression of a puromycin resistance transgene. The enzyme puromycin acetyl transferase (pac) or PuroR inactivates puromycin by N-acetylating its reactive amino group. Puromycin acetylation by pac requires the central metabolite and acetyl group donor acetyl-CoA as a substrate. We found that puromycin treatment exacerbates sensitivity of cancer cells to knockdown of pantothenate kinases, the proteins that catalyze the rate-limiting step of de novo coenzyme A production in cells. Mechanistically, we found that ablation of PANKs together with puromycin depletes acetyl-CoA levels, in a manner modulated by the dose of puromycin. Our findings provide a note of caution and context in the use of puromycin for metabolism research in that interference with the major acyl donor used for inactivating biotransformation may exacerbate toxicity under selection. Broadly, our findings also invite studies to explore how targeting CoA and acetyl-CoA synthesis may be exploited to enhance cytotoxic effects of cancer drugs that undergo acetylation.

## Introduction

Puromycin, an aminonucleoside antibiotic, inhibits protein synthesis by prokaryotic (70S) and eukaryotic (80 S) ribosomes (1). Puromycin is a close structural analog of 3’ end of aminoacylated tRNA (aa-tRNA) and constitutes a modified adenosine base which is linked covalently to a tyrosine amino acid by a peptide bond (2). Puromycin is incorporated into a nascent polypeptide chain on its C-terminus via a peptide bond (3). The presence of a stable peptide bond on puromycin (unlike a labile ester bond in an aa-tRNA) prevents further elongation of polypeptide chain resulting in premature termination of protein synthesis (2,3). Puromycin and its derivatives (such as radiolabeled (4,5), fluorescent (6,7), biotinylated (8,9), and photoactivable (10,11)), have been successfully used for applications including measurement of protein synthesis rate as well as visualization, and isolation of translating ribosomes and their products in cell-free systems, intact cells, and organisms (2). Additionally, puromycin derivatives have also been investigated for their promising potential in clinical applications as molecular probes for imaging (12,13), and as recent studies have highlighted as chemotherapeutics (14,15) and pro-drugs for targeted anti-cancer therapies (16,17). Puromycin is also a commonly used selection antibiotic in genetically engineered or transformed cells due to its desirable physical properties including water solubility well above the concentrations toxic to eukaryotic cells (1,2,18). Successfully transformed cells gain resistance to puromycin by inactivating biotransformation through expression of a transgene coding for puromycin N-acetyltransferase (pac) or PuroR, which is a bacterial enzyme found in *Streptomyces alboniger* (**Figure 1A**) (1,18,19). pac inactivates puromycin through acetylation of a nitrogen on the free amino group, catalyzing the acetyl-transfer from the central metabolite and acyl-donor acetyl-Coenzyme A (Acetyl-CoA) (**Figure 1A**) (1,18,19).

**Figure 1:**
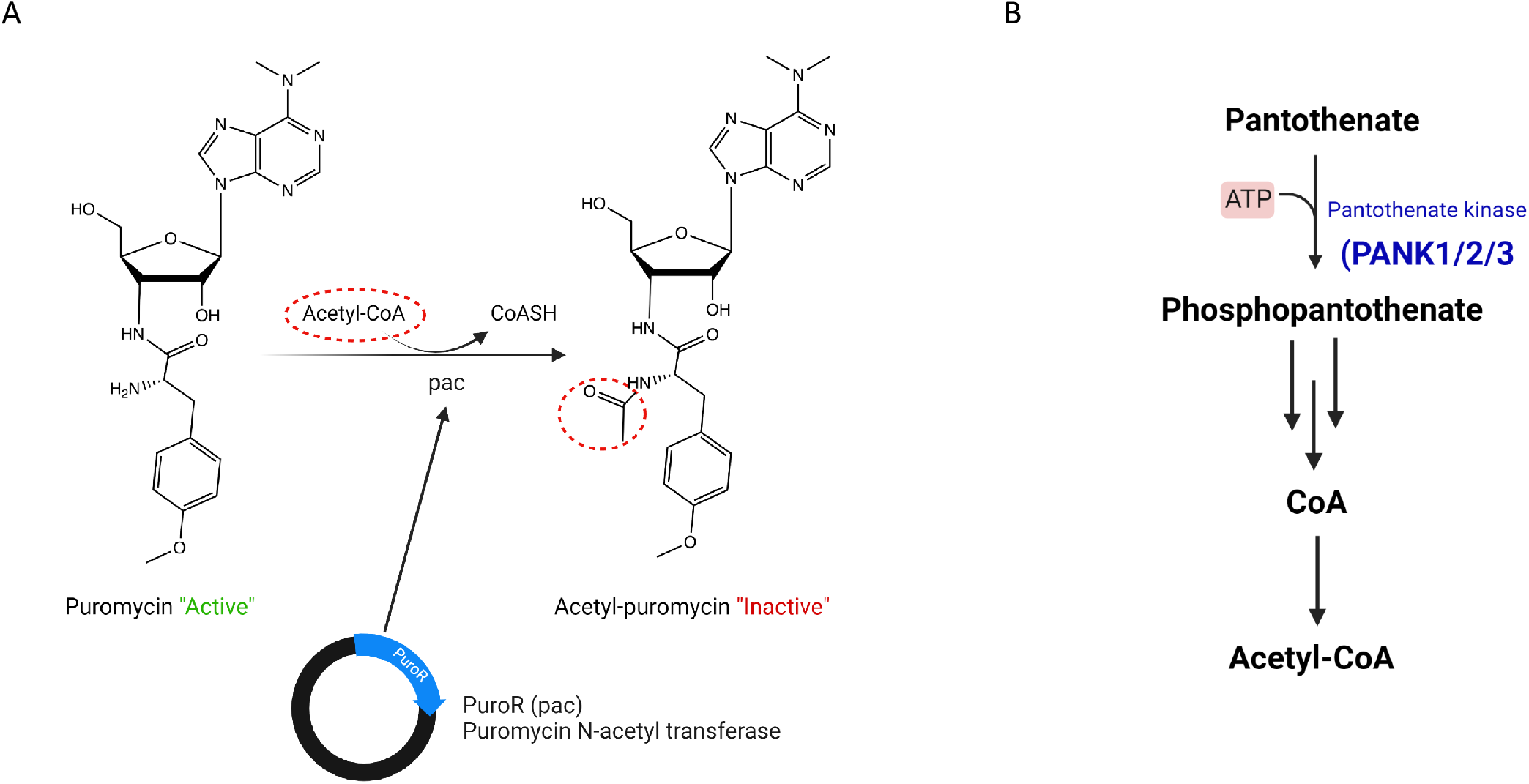
Puromycin acetyl-transferase inactivates puromycin via acetylation and consumes acetyl-CoA as a substrate. **A.** Puromycin, an aminonucleoside antibiotic, inhibits protein synthesis in prokaryotes as well as eukaryotes. As a tyrosyl-tRNA mimetic, puromycin forms a covalent peptide bond with the nascent polypeptide on ribosomes and prevents elongation and ultimately premature termination of polypeptide chain. Puromycin N-acetyl transferase inactivates puromycin via acetylation of its free amino group and uses acetyl-CoA as a substrate. PuroR or pac expression is routinely used as a selection marker to generate genetically engineered or transformed cells. **B.** Schematic showing the biosynthesis of CoA and acetyl-CoA. De-novo CoA biosynthesis in cells occurs with pantothenate (vitamin B_5_) as a substrate, which is converted to CoA in a series of biochemical reactions, the first and rate limiting reaction being controlled by pantothenate kinases (PANK1,PANK2 and PANK3). Acetyl-CoA is a major acyl carrier that sits at the intersection of central carbon metabolism in cells, and donates two carbon units to the TCA cycle, lipid and sterol biosynthesis, and participates in acetylation of proteins and metabolites.

Acetyl-CoA is a prototypical member of the evolutionarily conserved acyl-carriers that form a thioester bond to an acyl-chain with the CoA backbone assembled from pantothenate, cysteine, and adenosine (20). Acetyl-CoA is used in acetylation reactions of endogenous metabolites, xenobiotics, and proteins, as well as to provide 2 carbon units to the TCA cycle, de novo lipogenesis, and sterol synthesis (20,21). In prokaryotes as well as higher eukaryotes, de-novo CoA synthesis occurs from pantothenate, with the first and rate limiting step catalyzed by the pantothenate kinase (PANK) family of enzymes (**Figure 1B**) (22,23). Three PANK enzymes catalyze the conversion of pantothenate to 4’-phosphopantothenate (PANK1, 2, and 3), with PANK4 retaining phosphatase activity that is suggested to be critical in constraining cellular CoA concentration (22–25). Our investigations on CoA biosynthesis as a potential targetable vulnerability in cancers led us to assess whether genetic manipulation of the pantothenate kinases PANK 1, 2 and 3, as the key regulatory enzymes of de-novo CoA biosynthesis in prokaryotes and eukaryotes, could impact cancer cell viability.

An incidental observation in our ongoing studies was that ablation of pantothenate kinases yielded a significant exacerbation in cancer cell killing if the cells were concurrently treated with puromycin. Mechanistically, we found that puromycin administration together with pantothenate kinase ablation results in depletion of acetyl-CoA levels. This decrease in acetyl-CoA concentration exhibited a dose-response relationship with puromycin. Our data suggests that puromycin inactivation by pac in transformed cells can impact acetyl-CoA levels. Thus, we postulate that puromycin and pac expression may offer a tractable way to experimentally deplete cellular acetyl-CoA levels but this same phenomenon may confound the effect of manipulating cellular pathways that involve CoA and acetyl-CoA in some screening designs. Similarly, this effect of puromycin may suggests a strategy to potentiate the efficacy of anticancer drugs when used in conjunction with therapies that impinge on acetyl-CoA metabolism in cells.

## Methods

### Cell culture

The cell lines used in this work were HAP1(CVCL_YO91, leukemia) and HEK293T (CVCL_0063). The cell lines were authenticated at the MD Anderson Cytogenetics and Cell Authentication Core. HAP1 Wild Type (WT) and HAP1 *PANK1* knockout (KO) and HAP1 *PANK3* KO cells were commercially purchased from Horizon Discovery and HEK 293 T cells were purchased from ATCC. The cells were maintained in RPMI medium with 2mM glutamine (Cellgro/Corning #10-040-CV) (HAP1) or DMEM (Gibco™ #11995073) (HEK293T cells) supplemented with 10%FBS (Gibco/Life Technologies #16140-071) and 1% pen-strep (Gibco/Life Technologies#15140-122).

### Generation of stable dox inducible PANK1 and PANK3 shRNA cell lines

Stable doxycycline inducible shRNA expressing cells were generated using the third-generation lentivirus packaging system. A standard protocol was followed to generate recombinant lentiviral particles via transient transfection of HEK 293T cells. SMARTvector Inducible Lentiviral shRNA encoding a doxycycline inducible shRNA, puromycin resistance marker and green fluorescent protein (GFP) was purchased from Horizon Discovery. Briefly: 10 μg of the SMARTvector shRNA plasmid was mixed with 15 μg of packaging vectors (5 μg of pMd2g, 5 μg of pREV, 5 μg of p8.74) in OptiMEM. The plasmid mix were added to the OptiMEM + lipofectamine 2000 solution and transferred on to 293 T cells in 10 cm tissue culture dishes. The transfection media was replaced with fresh DMEM FBS 10% medium after 12 hours. For the next 48 hours since the first media change, viral supernatants were collected, and fresh medium was added to the HEK 293T cells every 24 hours. The viral supernatant was centrifuged at 500 g to eliminate cellular debris, and the virus supernatant was sterile filtered through a 0.45 μm filter. The target cells (HAP1) were transduced with the filtered viral particles by mixing the filtered viral supernatant with an equal volume of fresh medium and polybrene (8 μg/ml). After 24 hours, the media on the transduced cells was replaced with fresh media and 48 hours later, 2 μg /ml puromycin was used for selection. The transduced cells were maintained in 2 μg /ml puromycin containing RPMI medium.

### Crystal Violet Assay

Crystal Violet assay was used to determine the cell viability. Briefly, 2,000 cells per well were plates in 96-well plates in RPMI media. After incubating for 24 hours, cells treated with doxycycline with and without 2 μg/ml puromycin. After 4-5 days of drug treatment, cells were washed with phosphate-buffered saline (PBS) and fixed in 10% formalin. After fixation, cells were washed and airdried and crystal violet solution (0.05%) was used to stain the fixed cells. Following the removal of crystal violet and washing and drying of the plates, the dye was extracted with 10% acetic acid. Cell viability was quantified by spectrophotometric absorption of dye at 590 nm using a microplate reader.

### Western Blot

Cells were grown in 6 well plates and treated with 100 ng/ul doxycycline in RPMI medium. After 72 hours of doxycycline treatment, cell lysates were harvested by first washing the cells twice with ice cold phosphate-buffered saline (PBS) and then adding ice-cold RIPA buffer with protease (Complete™ mini, Roche#11836153001) and phosphatase inhibitors (PhosSTOP, Roche, #5892970001). The cell lysates were then sonicated.

Subcellular fractionation was performed on the HAP1 *PANK1* KO cells stably expressing the shRNAs following the manufacturer’s protocol (ThermoFisher #78840). Briefly, the cells were trypsinized, and ice-cold cytoplasmic extraction buffer was added on the trypsinized cells after washing with PBS. The cells were incubated in ice for 10 minutes with gentle mixing and supernatant was collected by centrifugation at 500g for 5 minutes.

Protein concentration was determined using the BCA assay (ThermoFisher, #23227). Nu-PAGE SDS-PAGE (4–12% gradient) was used to separate the proteins, which were then transferred onto nitrocellulose membranes using the semi-dry method (TransBlot turbo). Ponceau S staining was done to verify an effective transfer of proteins on the. The membranes were blocked in 5% non-fat dry milk in tris-buffered saline (TBS) with 0.1% Tween 20 (TBST) was. Primary antibodies were prepared in 5% milk and the membranes were incubated overnight at 4°C with gentle rocking. Next day, the membranes were washed 3x for 5 min with 1XTBST and incubated in HRP tagged secondary antibody (1:5000) for an hour with gentle rocking and then washed 3x for 5 min with TBST. ECL substrate (Thermo Fisher #32106) or ThermoScientific SuperSignal West Femto (#34096) was used as a substrate for HRP and the membranes were exposed on X-ray films in a dark room with different exposure times. The antibodies used in this study are as follows: PANK1 (CST #23887) PANK3 (MDA-299-62A) and Vinculin (CST #13901).

### CoA extraction and measurements

Acyl-CoAs were quantified by liquid chromatography-high resolution mass spectrometry as previously published (26). Briefly, cells were collected in 1 mL 4°C 10% (w/v) trichloroacetic acid, then frozen, and shipped for analysis. 50 μL of ^13^C_3_,^15^N_1_ acyl-CoA mix prepared as previously published from ^13^C_3_,^15^N_1_ pantothenate in yeast culture was added to each sample as an internal standard (27). Samples were mixed, sonicated with a probe tip sonicator, centrifuged at 17,000 x g 4°C for 10 minutes, then extracted by Oasis HLB solid phase extraction cartridges equilibrated with 1 mL of methanol, 1 mL of water, the sample, washed with 1 mL of water, and eluted with 1 mL of 25 mM ammonium acetate in methanol. The eluent was evaporated to dryness under nitrogen gas and samples were resuspended in 50 μL 5% 5-sulfosalicylic acid (w/v) in water. 10 μL of each sample was injected for analysis on an Ultimate 3000 UHPLC using a Waters HSS T3 2.1 x 150 mm 3.5 μm column on a gradient of 5 mM ammonium acetate in water to 5 mM ammonium acetate in 95:5 acetonitrile: water (v/v) with a wash solvent of 80:20:0.1 acetonitrile: water: formic acid coupled to a Q Exactive Plus mass spectrometer (Thermo Scientific). Each acyl-CoA was quantified by the [M-507+H]+ product ion with a 5 ppm window with Tracefinder 4.1 (Thermo Scientific). Analysts were blinded to sample identity during sample preparation and analysis.

## Results

### Puromycin significantly impairs the viability of PANK1 and PANK3 ablated HAP1 cells expressing PuroR

To interrogate if pantothenate kinases are redundant in cancer cells and whether these proteins (PANK1, PANK2 and PANK3) exhibit a synthetic lethal relationship, we engineered cancer cells that stably express doxycycline inducible shRNA against PANK1 or PANK3 proteins and selected the transformed cells by puromycin treatment. We employed commercially purchased *PANK1* or *PANK3* knockout HAP1 cells, and modulated PANK3 or PANK1 expression respectively, by doxycycline administration. HAP1 cells are near-haploid cancer cells derived from Chronic Myelogenous Leukemia. We performed crystal-violet based cell viability assay to determine the effect of genetic knockdown of PANK3 or PANK1 on the viability of *PANK1* or *PANK3* knockout cells (**Figure 2**). We found that knockdown of PANK3 or PANK1 modestly affects the viability of *PANK1* or *PANK3* KO cells compared to the non-targeting control (**Figure 2E-F, 2I-J**). However, we found that despite these cells being resistant to puromycin, because of PuroR or pac expression, administration of doxycycline in combination with puromycin significantly impaired the viability of these cells (**Figure 2G-H, K-L**). This indicated that puromycin’s mechanism of action potentially converges on de-novo CoA biosynthesis, which is regulated by PANKs, or associated pathways such as acyl CoAs, abrogation of which synergistically impaired the viability of HAP1 cells.

**Figure 2:**
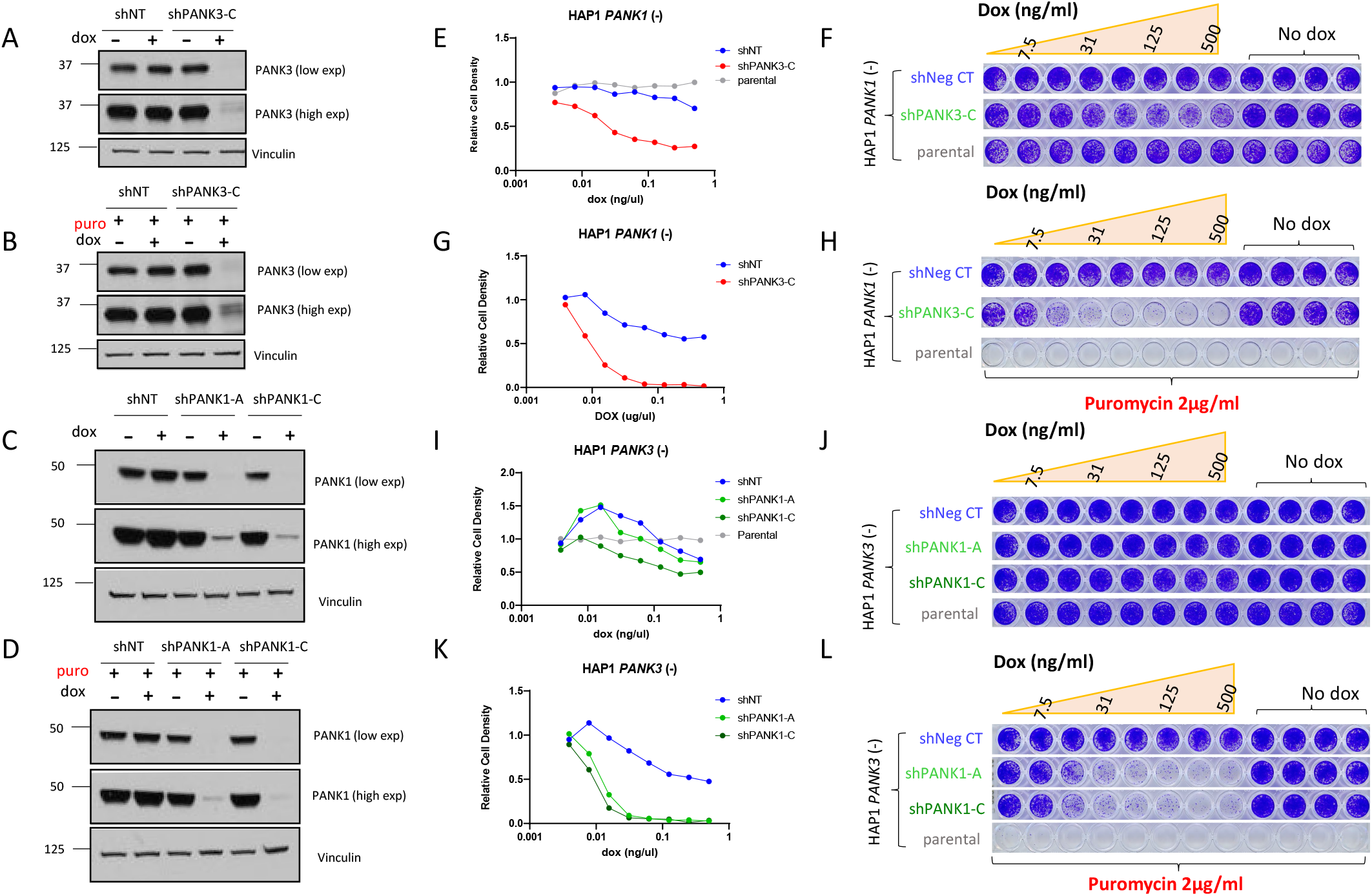
Puromycin exacerbates the toxicity of PANK ablation. (**A-D**)Western blot showing the knockdown of PANK3 or PANK1 protein in HAP1 *PANK1* KO or *PANK3* KO cells respectively with or without doxycycline (100 ng/ml) and puromycin (2 μg/ml). (**E-L**). Dose response curves of HAP1 *PANK1*(-) stably expressing doxycycline inducible PANK3 shRNA or HAP1 *PANK3*(-) cells stably expressing PANK1 shRNA treated with different concentrations of doxycycline, with or without 2 μg/ml puromycin. Cells were seeded at a density of 2000 cells per well in 96 wells plates in RPMI medium and treated with a serial dilution of doxycycline starting at 500 ng/ml. On day 4, cells were fixed with 10% formalin and crystal violet staining was performed to measure the terminal cell density and assess the effect of PANK knockdown in puromycin free or puromycin added medium. Puromycin was found to significantly exacerbate the toxicity of PANK knockdown in HAP1 cells.

### Puromycin exacerbates acetyl-CoA depletion in PANK ablated HAP1 cells expressing PuroR

The mechanism by which puromycin is inactivated in the cells expressing PuroR has been previously reported (18). Puromycin N-acetyl transferase inactivates puromycin through the transfer of an acetyl group on its reactive amino group and prevents puromycin from forming a peptide bond with a nascent polypeptide and inhibiting protein elongation (**Figure 1A**). PuroR mediated acetylation of puromycin requires acetyl-CoA as a substrate. We surmised that puromycin inactivation by PuroR affects cellular acetyl-CoA/CoA pools, and sensitizes cells to inhibition of related pathways, including CoA biosynthesis which is regulated by the PANKs.

To elucidate the mechanism behind the synergistic interaction between puromycin and PANK ablation, we profiled CoA and acyl-CoAs in HAP1 *PANK3* KO cells stably expressing shNontarget or shPANK1 with or without doxycycline and puromycin treatment (**Figure 3**). We observed that knockdown of PANK1 in *PANK3* KO HAP1 cells led to a decrease in CoA and acetyl CoA, only in combination with puromycin (**Figure 3D-F**). More importantly, we found that the degree of depletion of acetyl-CoA in these cells was dependent on the dose of puromycin (**Figure 3D**). This indicates that impinging on CoA biosynthesis by PANK ablation and depletion of acetyl-CoA by puromycin together disrupt cellular CoA and acetyl-CoA pools, leading to enhanced cancer cell death.

**Figure 3.**
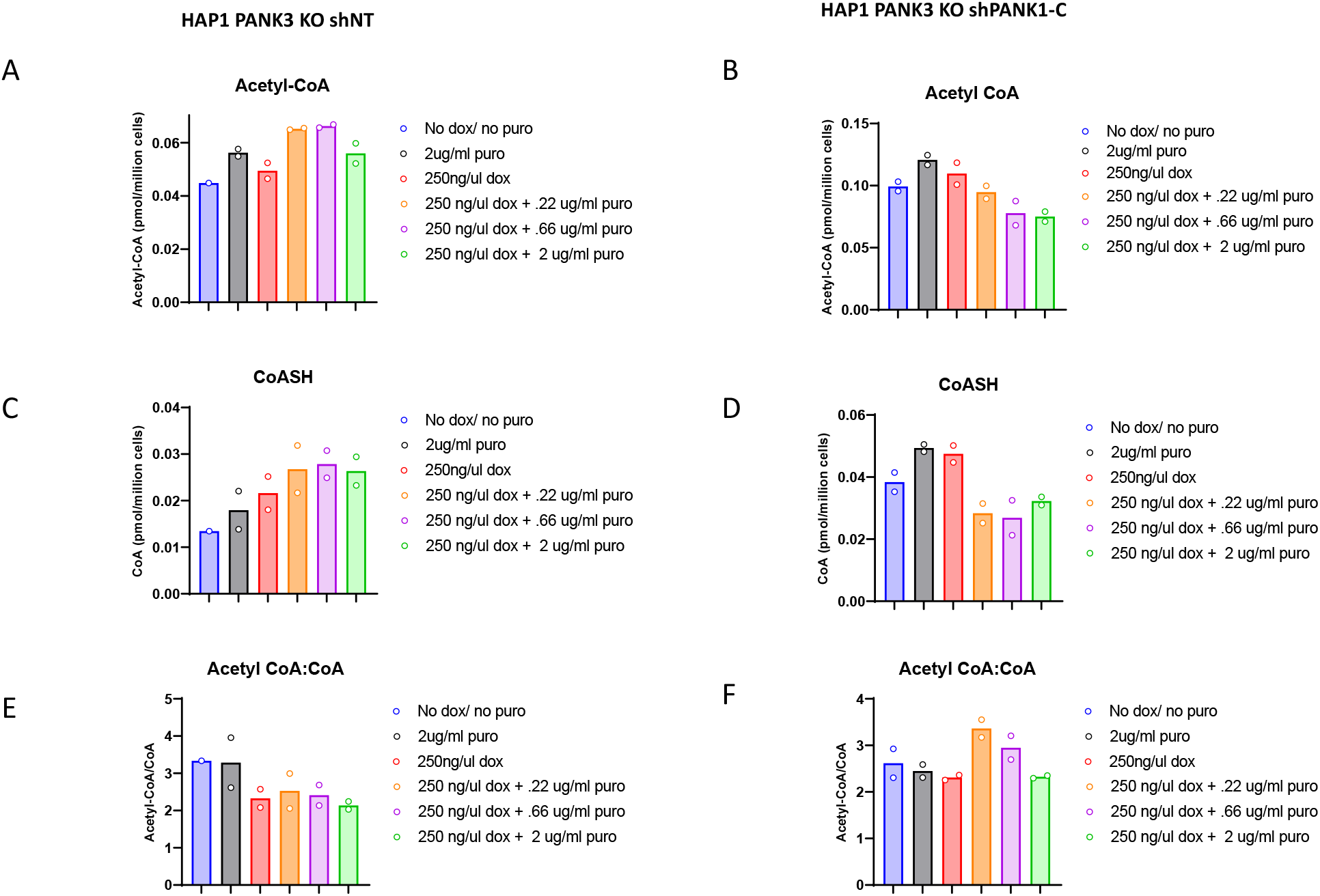
Puromycin exacerbates depletion of CoA and acetyl CoA in PANK ablated cells. HAP1 *PANK3* KO cells stably expressing doxycycline inducible shNontarget(shNT) or shPANK1 were treated with medium containing 250 ng/ml doxycycline with a serial dilution of puromycin for three days and assessed for the changes in acetyl CoA and CoA levels. Acetyl CoA, CoA and Acetyl-CoA:CoA ratio for each treatment conditions are shown. Puromycin administration exacerbates a dose dependent depletion of acetyl CoA specifically in shPANK1 expressing cells.

**Figure 4.**
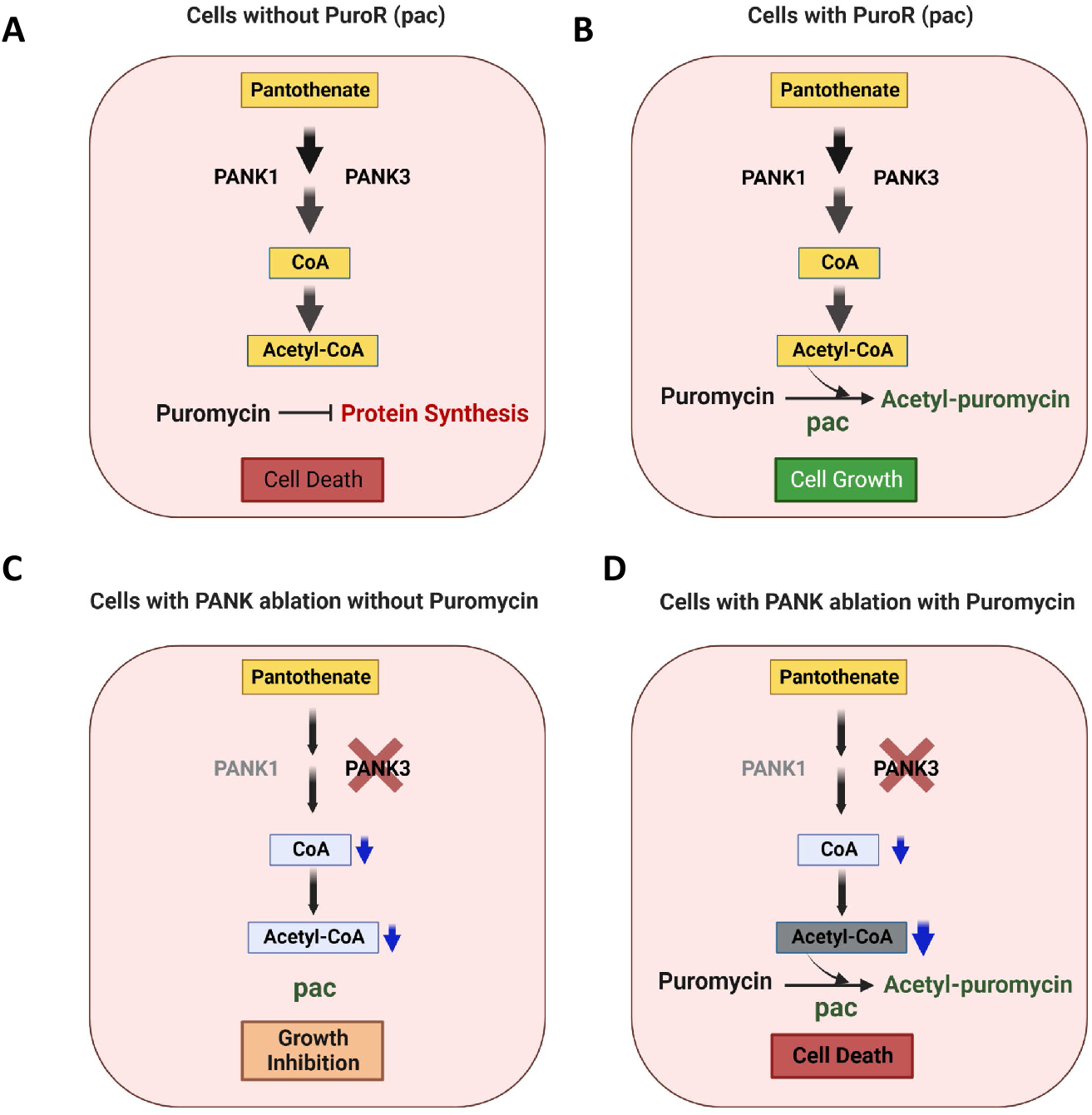
Graphical abstract: A. Puromycin inhibits protein synthesis in cells that do not express PuroR, resulting in their death. B. PuroR acetylates puromycin, inactivating it, which prevents puromycin’s toxic effects on cells. C. PANK ablated cells have lower acetyl-CoA levels, which leads to growth inhibition. D. PuroR expressing cells that are PANK ablated are significantly sensitized to puromycin, which exacerbates cell killing through depletion of acetyl-CoA beyond toxic threshold.

## Discussion

In mammalian toxicology, N-acetyl-transferases (NAT) 1 and 2 are the predominant enzymes for xenobiotic detoxification through N-acetylation biotransformation of aromatic amines (28,29). The NATs have broad substrate specificity but less affinity for their xenobiotic substrates, with primary toxicological examples of substrates including aromatic amines (30). To our knowledge, no reports of substrate limitation exist for N-acetyltransferase mediated phase II metabolism. In contrast to NATs, pac has high affinity for puromycin. This makes pac desirable as a drug resistance enzyme especially for selection of resistance in molecular biology (1,18).

Our study is limited by the preliminary nature of this report and that acetyl-CoA is almost certainly a necessary metabolite for cell viability. Thus, some genetic manipulations of PANKs (a triple knockout of PANK1, 2, 3) are likely not viable, nor would a depletion of acetyl-CoA beyond some thresholds be viable. Additionally, acyl-CoAs exist in separate sub-cellular pools within the eukaryotic cell, and puromycin/pac may selectively deplete one or more pool while leaving others relatively intact. Future work will be needed to examine the effect of the inhibition of acetyl-CoA generating enzymes such as ATP-citrate lyase (ACLY), Acetyl-CoA Synthetase 1 (ACSS1), and ACSS2 on this phenomenon.

These limitations in the scope of the future work also should prompt examination of screening designs based on puromycin selection. Mutations in or selection based on pathways, enzymes, or conditions that modulate acetyl-CoA may result in negative or positive selection due to the presence of puromycin. Selection by blasticidin (which relies on a Cytochrome P450 mediated deamidation for inactivation) would not be expected to result in the same phenomenon.

This phenomenon of interaction between acetyl-CoA and puromycin resistance also offers opportunities for experimental design and cancer pharmacology. The selective reduction in acetyl-CoA by pac catalyzed N-acetyl transfer to puromycin could be taken advantage of for the purposes of modulating the cellular acetyl-CoA pool. To our knowledge, loss of inactivating biotransformation due to depletion of the acetyl-CoA substrate has not been previously observed. This indicates that in contexts where PANK loss may occur such as co-deletion of *PANK1* with *PTEN* loss, selective cellular toxicity may be more readily achievable using pharmacological agents that decrease CoA availability and require detoxification by acetylation.

